# Dynamic representation of sound locations during task engagement in marmoset auditory cortex

**DOI:** 10.1101/2025.08.14.669832

**Authors:** Chenggang Chen, Evan D. Remington, Xiaoqin Wang

## Abstract

In auditory cortex, neural responses to stimuli inside receptive fields (RFs) can be further facilitated by behavioral demands, such as attending to a spatial location. It is less clear how off-RF stimuli modulate neural responses and contribute to behavioral tasks. Our recent study revealed a particular form of location-specific facilitation evoked by repeated stimulation from an off-RF location, suggesting behavioral modulation of spatial RFs. To further explore this question, we trained marmosets to attend to sound locations that were either inside or outside the RFs of auditory cortical neurons. The majority of neurons showed increased firing rates at target locations inside their RFs. Interestingly, this increase also occurred outside the RFs, sometimes exceeding the responses at the RF center during passive listening. These task-related off-RF facilitation were much more common in the caudal area than in the rostral area and the primary auditory cortex. A normalization model reproduced the off-RF facilitation using widespread suppression. The model’s prediction was confirmed by experimental observations of widespread reductions in firing rate and hyperpolarized membrane potentials for off-RF stimuli. These results suggest that behavioral task demands recruit a broader range of neurons than those that are responsive to a target sound in the passive state.

## Introduction

It is well known that response properties of neurons in the visual (Ruff and Cohen, 2019; Ghosh and Maunsell, 2022; Jonikaitis et al., 2025; Srinivasan et al., 2025), somatosensory (Mima et al., 1998; Steinmetz et al., 2000; Gomez-Ramirez et al., 2016), and auditory (Otazu et al., 2009; Mesgarani and Chang, 2012; Niwa et al., 2012; McGinley et al., 2015; Heller et al., 2024; Lu et al., 2025) cortex can change rapidly to match specific task demands. In the visual cortex, this phenomenon has been studied extensively for visual location (Fig. 1A), for which attending a behaviorally relevant locations inside receptive fields (RFs) increase neural activity (Luck et al., 1997; McAdams and Maunsell, 1999; Treue and Martinez-Trujillo, 1999; Lee et al., 2007; Mitchell et al., 2007; Anderson et al., 2011; Yao et al., 2016; Speed et al., 2020) and reduce the inter-neuronal correlations (Cohen and Maunsell, 2009; Mitchell et al., 2009; Nandy et al., 2017). Similarly, attending to different locations of somatosensory stimuli inside the RFs of the finger or whisker also facilitates neural responses (Hsiao et al., 1993; Ramamurthy et al., 2025). In auditory cortex, extensive work on frequency tuning (Fig. 1B) shows sharpened RFs (Ohl & Scheich 1996; Okamoto et al. 2007, 2009; Neelon et al. 2011; O’Connell et al. 2014; Xin et al. 2019) and mainly enhanced, occasionally suppressed, responses at target frequencies within the RFs of single neurons or the majority of simultaneously recorded neurons (Fritz et al. 2003, 2007; Atiani et al. 2009; David et al. 2012; Kato et al. 2015; Kuchibhotla et al. 2017; Francis et al. 2018; Insanally et al. 2024).

**Figure 1.**
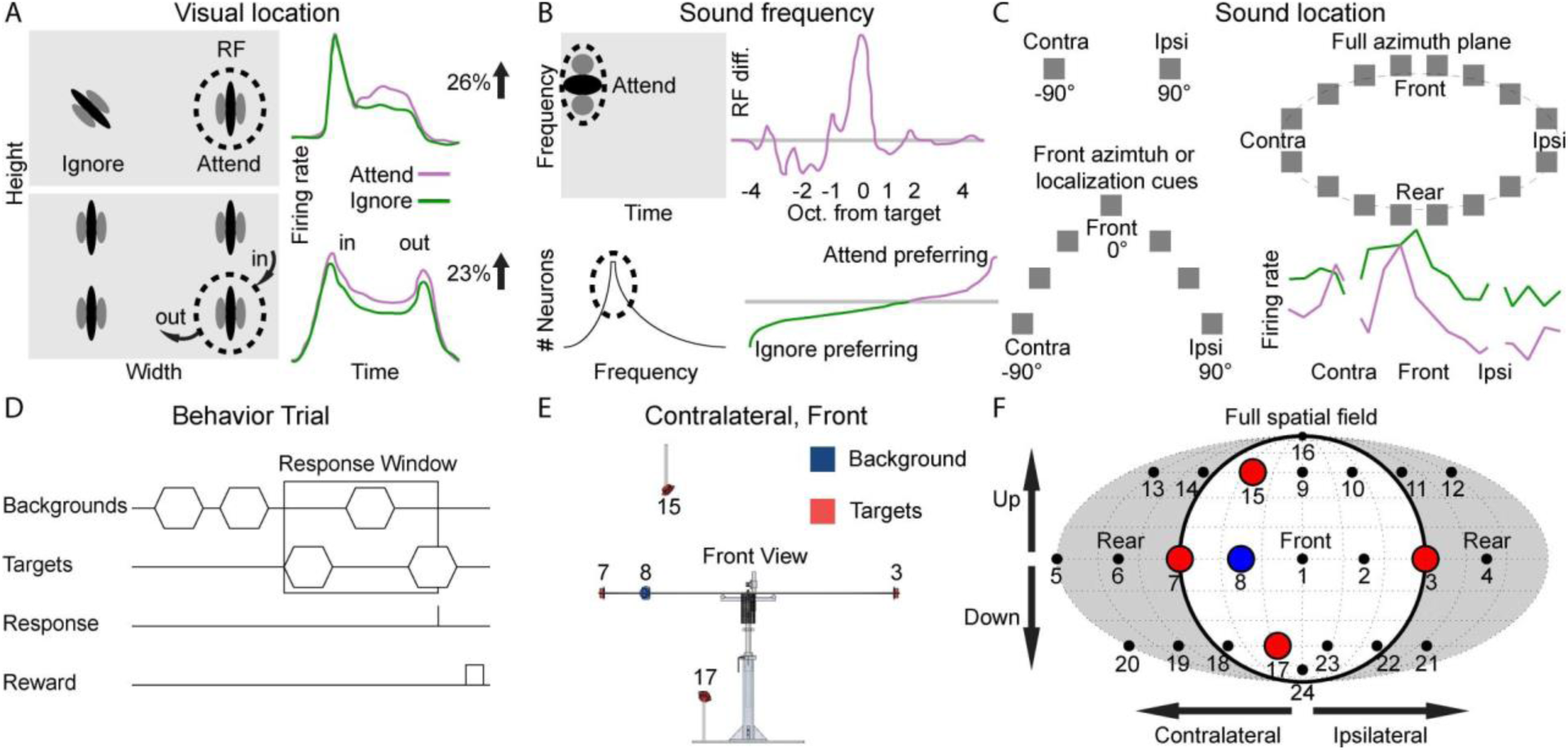
Comparison of behavioral tasks in previous and current studies. (A) Two visual spatial attention tasks and firing rate changes. Top, two Gabor stimuli were presented inside and outside of the receptive field (RF). The animal is trained to either attend to or ignore the stimulus inside the RF. Attending to the stimulus inside the RF increases the firing rate (modified from Cohen and Maunsell, 2009). Bottom, all four stimuli moved along randomized trajectories that brought one stimulus into the RF. All stimuli then paused for 1000 ms, and one stimulus moved out of the RF (modified from Mitchell et al., 2007). (B) Two auditory sound frequency attention tasks and firing rate changes. Top-left, the spectrotemporal response field (STRF) with one center excitatory (dark) and two inhibitory (gray) regions. The frequency of the target tone fell inside the STRF. Top-right, average spectral change in the STRF at all frequencies relative to the target frequency (unit: octave) (modified from Fritz et al., 2003). Bottom-left, best frequency histogram of all simultaneously recorded neurons, and the target was chosen inside the RF. Bottom right, histogram of neurons that showed enhanced and suppressed responses when attending the target frequency (modified from Kuchibhotla et al., 2017). (C) Subjects were required to attend to only contralateral or ipsilateral 90° (left-top), front azimuth or azimuth localization cues (left-bottom), or full azimuth plane (right-top). Average firing rate as functions of azimuth and ipsilateral and contralateral 90° were omitted from analysis (modified from Lee and Middlebrooks, 2011). (D) After a variable number of background stimuli, targets will begin alternating with the background stimuli. If a lick is registered within the preset number of alternations, a food reward is given. (E) Speaker layout for one of the four behavior conditions. The background location was 45° off midline, and the target locations were the most lateral positions (±90°; same in all conditions), and also 45° above and below the horizon, but in the same azimuthal quadrant as the background location. (F) Fournier map projection of a complete 24-speaker spatial array with speakers from (E) highlighted.

A common procedure for a sensory attention task is to present a target stimulus that falls inside or near the RFs of recorded neurons. Visual or somatosensory location and auditory frequency are faithfully relayed from the sensory receptors to the cortex, with topographic maps preserved across all the stations along the ascending pathways (Arcaro and Livingstone, 2024). A stimulus far from the RF is not supposed to evoke neural response. In contrast, the sound location information is computed from the sensory inputs from two ears by the auditory system (Grothe et al., 2010; Yin et al., 2019) and is then used in a variety of behavioral contexts beyond sound localization (Shinn-Cunningham, 2008; Bizley and Cohen, 2013; van der Heijden et al., 2019; Francl and McDermott, 2022), such as sound segregation in a cocktail-party (McDermott, 2009). We recently showed that when a sound stimulus was repeatedly presented from a particular location outside the RF of a neuron, the neuron could temporally increase its responses to this location, suggesting facilitation or attention-based mechanisms (Chen et al., 2025). To fully understand such mechanisms, one needs to first sample the entire auditory spatial field beyond the azimuth and horizonal axes to determine the extent of the RF of a neuron. However, sampling spatial locations that fall over a three-dimensional sphere (not considering distance) is technically challenging in neurophysiological recordings.

Auditory cortex is essential for many behaviors involving sound localization in mammals (Jenkins and Masterton, 1982; Bizley et al., 2007; Lomber and Malhotra, 2008; Bajo et al., 2010, 2019; Nodal et al., 2010; Wood et al., 2017; Town et al., 2023). Neural responses in the auditory cortex are also modulated by spatial attention. Studies in humans using non-invasive imaging methods (Zatorre et al., 1999; Petkov et al., 2004; Zimmer and Macaluso, 2005; Ahveninen et al., 2006; Degerman et al., 2006; Hill and Miller, 2010; Ding and Simon, 2012; Lee et al., 2013; Kong et al., 2014; Higgins et al., 2017; Wang et al., 2022) and surface electrode (Patel et al., 2022) all found enhanced responses to when attention is directed to preferred sound locations, except one study which shows similar responses to preferred locations but decreased responses at nonpreferred locations (van der Heijden et al., 2018). Animal studies, which have single-neuron resolution, have shown varied effects of task engagement on responses in the auditory cortex. These effects include increased firing rates to sounds played to an attended ear in a selective listening task (Benson and Hienz, 1978), increased firing rates to arbitrary locations in a localization task (Benson et al., 1981), and general firing rate increases in an interaural phase discrimination task (Scott et al., 2007). In contrast, in an elevation discrimination task, responses to preferred locations are consistent, but to non-preferred locations decreased, resulting in narrower tuning widths (Lee and Middlebrooks, 2011, 2013). In addition, for the spatially untuned neurons in a free navigation task, half of them increased and the other half decreased responses to the target speaker (Amaro et al., 2021). Together, previous studies suggest that attending to preferred sound locations increases neural activity, but the effect could be facilitative or suppressive at nonpreferred locations.

All of the aforementioned studies measured responses to a subset of spatial locations (Fig. 1C), including contralateral and ipsilateral (Benson and Hienz, 1978; Zatorre et al., 1999; Petkov et al., 2004; Degerman et al., 2006; Ding and Simon, 2012; Lee et al., 2013; Patel et al., 2022), front azimuth (Benson et al., 1981; Ahveninen et al., 2006; Hill and Miller, 2010; van der Heijden et al., 2018; Wang et al., 2022), azimuth localization cues (Zimmer and Macaluso, 2005; Scott et al., 2007; Kong et al., 2014; Higgins et al., 2017) or full azimuth plane (Lee and Middlebrooks, 2011, 2013; Amaro et al., 2021). The definition of preferred and nonpreferred locations based on a partial spatial field is inaccurate and could be misleading until the full spatial field (both azimuth and elevation) is sampled. However, little is known on how auditory cortex neurons represent the full spatial field (Middlebrooks and Pettigrew, 1981; Brugge et al., 1994, 1996; Schnupp et al., 2001; Mrsic-Flogel et al., 2005), let alone test it during behavioral engagement. For the first time, we mapped the neural representations of the full spatial fields in the auditory cortex of awake, passively listening subjects (Remington and Wang, 2019). This study shows that widespread distribution of preferred locations of auditory cortex neurons, which is not limited to the front space or azimuth plane. The importance of using a larger sampling space was further exemplified by our previous work showing that a masker sound from a nonpreferred location inhibited neural responses to a probe sound from a preferred location (Zhou and Wang, 2012, 2014), and our recent work showing neural facilitation to repetitive stimuli from nonpreferred locations outside a neuron’s spatial RF (Chen et al., 2025).

To address these questions, we recorded single-unit responses in the auditory cortex of marmosets (*Callithrix Jacchus*), an arboreal New World monkey, while they performed a spatial discrimination task (Chen et al., 2023). Unlike earlier auditory attention studies that sampled only a portion of space, we trained marmosets to attend to targets from the entire spatial field. Presenting full field stimuli enabled us to accurately identify each neuron’s spatial RF during a passive state, and then measuring task-related modulations in both preferred and non-preferred locations. We found a task-related facilitation effect for both preferred and non-preferred locations. Taken together with our recent findings (Chen et al., 2025), these results suggest that the classical, static view of cortical spatial RFs must be revisited. Dynamic spatial responses, in particular outside RFs, could support functions such as auditory streaming and the cocktail-party effect, and possibly account for the absence of an ordered cortical map of auditory space (Middlebrooks, 2021).

## Results

We recorded 208 single units from two marmosets while the animals either listened passively to broadband sounds played from speakers on a complete sphere or performed a sound location discrimination task that included a subset of those locations (Fig.1E, F). Firing rates in the behaving condition were compared to two other conditions, a “passive condition” in which we played sounds in a random order from the full 24-speaker array and used the half-maximum interpolated firing rates as the spatial receptive fields (RFs, Remington and Wang, 2019), and a “control condition” in which sounds were played only from target/background locations in the same pattern as the behavior condition and with roved sound level. The latter was used to control for the potential confounding effects of stimulus order (Chen et al., 2025). All analyses were conducted on stimuli presented in successful discrimination (“Hit”) trials with spontaneous firing rates subtracted.

### Attention increased the firing rates at target locations outside the RFs

Figure 2A shows the spatial RF, which was computed using the passive response to 24 randomly presented locations shown in Fig. 2B (blue dots). This unit has a complex RF (black curves), with responses distributed among contralateral bottom and rear locations. In the contralateral front configuration, all four targets and one background were outside the RFs. This unit displayed increased firing rates to two of four targets (#7 and #15), as well as the background (#8), in the behaving condition (Fig. 2C). In the ipsilateral, front configuration, we observed a consistent and even stronger firing rate increase at location #7 (Fig. 2D, bar).

**Figure 2.**
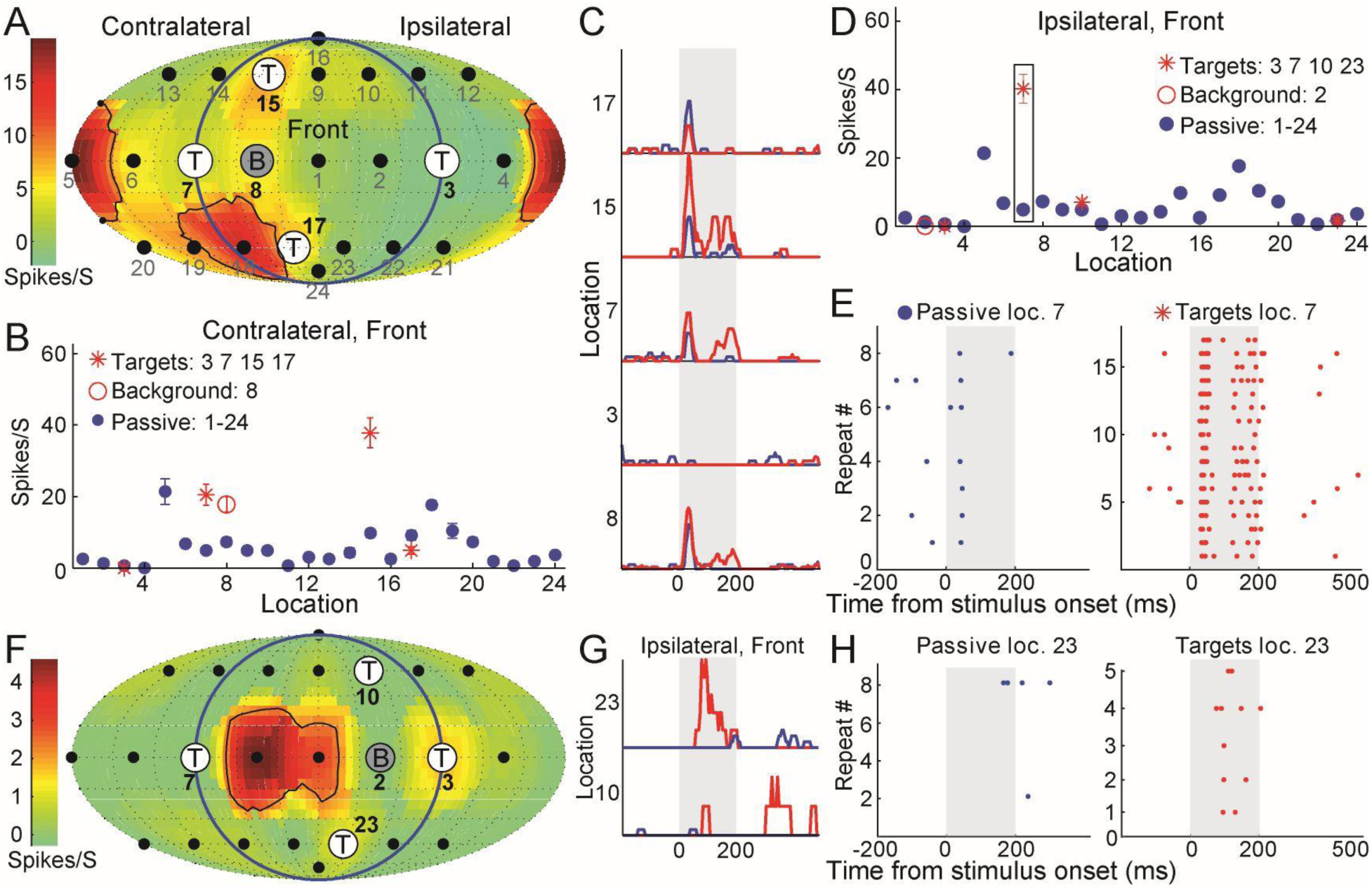
Firing rates increase at locations outside the RFs during spatial discrimination. (A) Passive spatial RF in one animal (M9X-74). Speaker locations, including target and background locations of the discrimination task configuration in Figure 1, were overlaid on the RF. (B) Comparison of firing rates measured in the passive (blue) and behaving (red) conditions. Error bars denote the standard error of the mean. (C) Peri-stimulus time histogram for four targets (#3, #7, #15, and #17) and one background (#8) location in each condition. The vertical bar indicates 200-ms sound stimuli. (D) Similar to (B) but with a different background (#2) and targets (#10 and #23). (E) Spike raster in passive (left, 8 trials) and targets (right, 17 trials) conditions from location #7 highlighted within the bar inside (D). (F) Passive spatial RF in the other animal (M71V-2522). The targets and background locations were the same as (D). (G) Peri-stimulus time histogram at two target locations. (H) Spike raster in passive (8 trials) and targets (right, 5 trials) conditions from location #23.

Furthermore, spike raster at the target but not the passive location showed highly reliable and consistent pattern among repeated stimulus presentation (Fig. 2E). In the contralateral, rear configuration, we observed firing rate increase at all target locations except #3 (Supplementary Fig. 1A). Figure 2F and Supplementary Fig. 1F shows a simple RF from another unit. This unit has a near-zero spontaneous firing rate and zero firing rate in 20 locations. However, its firing rates were strongly facilitated at target #23 (Fig. 2G). Importantly, this unit showed reliable spikes during all the five repeats of target (Fig. 2F). Compared with the target, we did not find reliable firing rate increase at the background (Supplementary Fig. 1B). Targets located inside the RFs can either had strongly increased (Supplementary Fig. 1C, #15) or zero (Supplementary Fig. 1E, #7) firing rate comparing to passive condition. All four targets located outside the RFs can increase firing rate (Supplementary Fig. 1D). Supplementary Fig. 1C-F also showed the firing rate changes under the control condition (blue dots and black lines), and we found it was similar to the passive condition but far different from the behaving condition. In summary, comparing targets with passive/control conditions, there was a majority of increased firing rate to one or more specific target locations.

### Quantification of attention modulations inside and outside the RFs

Figure 3A-C shows the population distribution of behaving vs. passive comparisons for each target and background location tested for each unit. Population analyses included neurons that displayed a driven firing rate (p<0.001 and minimum mean rate of one spike per stimulus presentation) to at least one location in at least one condition. Although in most studies spontaneous rates have not been observed to change significantly in behavior (Benson and Hienz, 1978; Otazu et al., 2009), several studies found increased spontaneous rates during behavior (Scott et al., 2007). In this sample population, there was a modest but significant increase in spontaneous firing rates when comparing hit trials to passive listening (p = 3.32 x 10^-4^, Wilcoxon signed-rank test). We quantified the difference using the modulation index (MI), which is a measure of response difference between two conditions (hits minus passive or control), scaled by the combined responses (hits add passive or control). MI ranges from -1 to 1, and a positive MI indicates the firing rate from one location was larger during hits (either at target or background locations) than during passive or control. All following analyses treat each location tested in each neuron as a separate data point.

**Figure 3.**
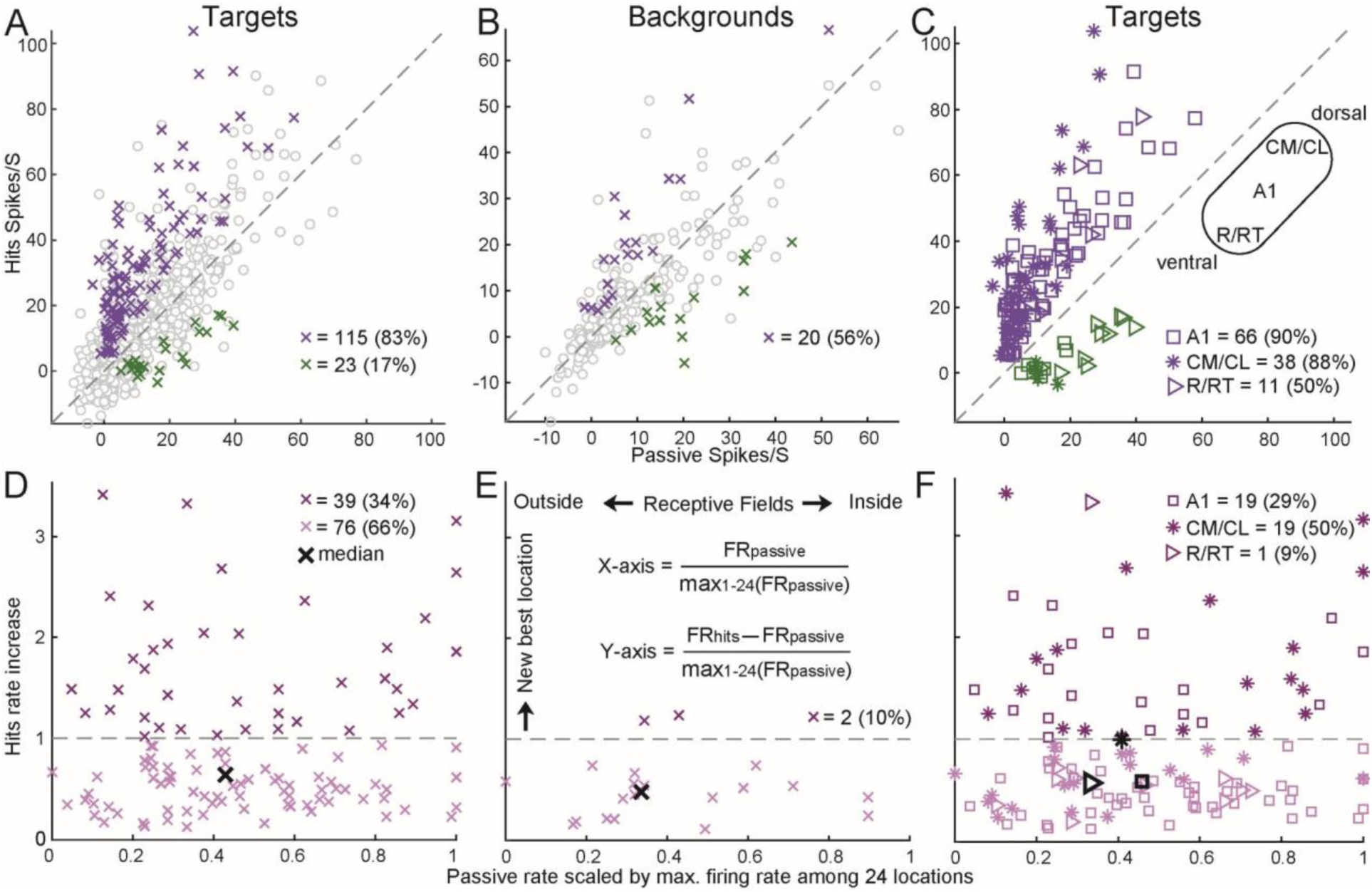
Attention increased firing rate and generated temporary RFs at target locations. (A) Comparisons of firing rates in the hit (i.e., successful behavioral choice) and passive conditions at target locations. All firing rates are spontaneous rates subtracted. Colored crosses represent significantly (p < 0.05) increased (pink) and decreased (green) firing rates at target locations. Gray circles represent the nonsignificant locations (576 (53%) above the diagonal). (B) Similar to (A) but at the background locations. There were 143 (49%) circles above the diagonal. (C) Same data as the colored crosses shown in (A), but with three different shapes (asterisk, square, and right-pointing triangle) to distinguish the three auditory cortical areas (A1, CM/CL, and R/RT). (D) Hits firing rate increases relative to passive firing rate (Y-axis) against passive firing rate (X-axis) for all increased data points (115 pink crosses in (A)), both scaled by the maximum firing rate among 24 locations in the passive condition. Median increase is 0.62 (Y-axis). (E) Similar to (D) but for background locations (median: 0.49). (F) Same data as (D) but distinguished A1, CM/CL, and R/RT (median: 0.56, 0.98, and 0.55).

For target locations that displayed behavior modulation during hit trials (Fig. 3A), we observed elevated firing rates in the majority of locations (83%, purple) during the behaving condition (Y-axis) compared with the passive condition (X-axis). The MI was 0.48 (median, hereafter) for all target and passive pairs. Of 208 units tested, 70 (79%) had at least one significantly increased firing rate, while 19 (21%) had at least one significantly decreased rate. In contrast, for background locations (also in hit trials, Fig. 3B), the effects were mixed: we observed both increased and decreased firing rates, with an MI of 0.21. There were 17 (53%) units that had at least one significantly increased firing rate to backgrounds, and 14 had at least one significantly decreased firing rate. In the control comparison, MI for targets of 0.50 (Supplementary Fig. 2A) was similar to the passive comparison; of 111 units tested, 37 and 2 had at least one significantly increased and decreased firing rate to targets, respectively. The control MI for backgrounds of 0.26 (Supplementary Fig. 2B) was larger than for the passive comparison but did not reach significance (p = 0.15); 16 and 5 units had at least one significantly increased and decreased firing rate to backgrounds, respectively. Therefore, consistent with many previous studies, attending to targets but not background locations increased neural activities. Importantly, as shown in the example neurons (Fig. 2 and Supplementary Fig. 1), the increase occurred for neurons even with a very low passive firing rate (near zero on the X-axis).

Many studies have shown that the distributions of spatial tuning properties vary between auditory areas along the rostral-caudal axis (Fig. 3C, inset), with neurons in caudal areas (CM/CL), on average, displaying higher selectivity for spatial locations than those in rostral areas (R/RT) and primary auditory cortex (A1) (Tian et al., 2001; Stecker et al., 2003; Woods et al., 2006; Zhou and Wang, 2012; Lee and Middlebrooks, 2013; Remington and Wang, 2019). However, studies that compared the effects of engagement in a sound localization task among three areas revealed conflicting findings. The fMRI studies found that spatial tunings in the human posterior auditory cortex are either task variant (Higgins et al., 2017) or invariant (van der Heijden et al., 2018). A neurophysiology study found stronger task modulation in the cat caudal area (Lee and Middlebrooks, 2013). We therefore asked whether any behavioral effects would differ quantitatively along the rostral-caudal axis in a primate species. Units were separated by area using frequency map gradient reversal points (see methods), and a significant effect of area on MI was observed when comparing behaving and passive conditions (Kruskal-Wallis ANOVA, p = 4 x 10^-4^; p = 0.03 for A1 vs. CM/CL; p = 0.02 for A1 vs. R/RT; p = 0.003 for CM/CL vs. R/RT, corrected for multiple comparisons), finding the largest values in CM/CL, and the smallest in R/RT (Fig. 3C). There were 36 (86%), 24 (83%), and 10 (56%) significantly increased units for A1, CM/CL, and R/RT. When this analysis was performed using behavior/control comparisons (Supplementary Fig. 2C), the trend was similar; however, differences were no longer significant (Kruskal-Wallis ANOVA, p = 0.1) except for the CM/CL to R/RT comparison. This lack of effect is likely partially due to the reduced statistical power of the behavior /control comparisons (n = 37 units compared with 86 for the behavior/passive comparison), but also due to the increased MI (0.42) in areas R/RT. This suggest that the stimulus order in the behaving/control condition may play a role in depressing firing rates, particularly in areas R/RT. Together, attention primarily increased the firing rates in A1 and CM/CL neurons but the effect was mixed in R/RT neurons.

### Attending to target locations shifted spatial response profile

Firing rate changes during sound localization tasks have been observed to be correlated with underlying spatial RFs (Benson and Hienz, 1978; Lee and Middlebrooks, 2011, 2013), but not in all cases (Benson et al., 1981). As we discussed in the introduction, all those studies measured firing rate to only a subset of spatial locations (Fig. 1C). The interpretation of these effects will benefit from a more complete sampling of spatial RFs, as evidenced by our recent studies that showed location-specific facilitation using a half spatial field (Chen et al., 2025). In this study, all the example units (Fig. 2 and Supplementary Fig. 1) and population analysis (Fig. 3A-C) showed attention increased firing rate at the nonpreferred target locations with weak or no firing rates. Here, we quantified the increases occurred along the passive response tuning curve, which is the passive firing rate scaled by the maximum passive firing rate among 24 locations (Fig. 3D-F). On the X-axis, a value of 1 indicates the target/background locations evoked the maximum responses from the center of RFs (e.g., #15 in Supplementary Fig. 1c was next to the center of RF). A value of 0.5 indicate the locations fell near the boundary of RFs (e.g., #17 and #15 in Fig. 2A). A value close to 0 indicate the locations fell outside the RFs (e.g., #3 in Fig. 2A). Among the 115 target locations that exhibited significantly increased responses during hits than passive locations (pink crosses, Fig. 3A), the median of scaled passive rate (0.43, black cross, Fig. 3D) was close to boundary of RFs (i.e., 0.5). Among the 20 background locations (Fig. 3B), the median of scaled passive rate was 0.33 (Fig. 3E). The difference of two values indicates that during the passive condition, the target locations which include both azimuth and elevation evoked larger responses than azimuth only background locations.

The Y-axis in Fig. 3D-F measured the magnitude of increase that was also scaled by the maximum firing rate at the center of RFs. We found that the increases in firing rate occurred throughout RFs, as indicated by the large number of increases that occurred for locations outside the RFs, which drove the neuron poorly in the passive condition. Increases did not occur at all locations in units with increased responses, and this effect was not due to the predominance of non-driven locations, which did not often show firing rate increases. Out of a minimum of 4 locations tested, the median number of significantly increased responses per significantly increased unit was 1 location, while the median number of driven locations in the same population of units was 3 (p = 4 x 10^-9^), indicating that increases did not occur simultaneously across RFs; rather effects were location specific. The horizontal dashed lines (value of 1) indicate that the magnitude of increase was larger than the firing rate at the RF center. Surprisingly, attending to a location generated a new best location (i.e., RF center) temporarily among 34% of the target but only 10% of the background locations.

We further distinguished three cortical areas for the targets using the same data shown in Fig. 3D. Although the percent of significantly increased locations were similar between A1 and CM/CL (Fig. 3C), the percent of locations above horizontal line was higher in CM/CL than A1 (Fig. 3F). The medians of scaled hit firing rate increases were similar between A1 and R/RT, but was much higher in the CM/CL (closer to 1). The difference between CM/CL versus A1 and R/RT was also significant (p = 0.0013, rank-sum). Together, attention generated new spatial responses outside RFs mainly in the CM/CL neurons.

### Modeling the dynamics of spatial responses with widespread suppression

What is the neural mechanism underlying this firing rate increase both inside and outside the spatial RFs? To model this auditory spatial attention effect, we choose the widely used divisive normalization model (Carandini and Heeger 2012). Normalization model was first developed to model visual responses to stimuli of different contrast (Heeger, 1992), and subsequently used for visual spatial attention (Reynolds et al., 1999; Ghose and Maunsell, 2008; Lee and Maunsell, 2009; Reynolds and Heeger, 2009), spectrotemporal contrast (Rabinowitz et al., 2011), and multisensory integration (Ohshiro et al., 2011). Importantly, some predictions of the normalization model have been validated experimentally (Herrmann et al., 2010; Lee and Maunsell, 2010; Ohshiro et al., 2017).

There are three fields in this model (Fig. 4A): stimulus (left), attention (top), and normalization (or suppression, bottom). On each simulated trial (or target location), the stimulus field is multiplied by the attention field and divided (i.e., normalized) by the suppressive field to generate the output firing rate. We modeled all three fields as two–dimensional Gaussian kernels within a 29 × 29 grid. The stimulus field was always centered and had a fixed standard deviation (σ = 2). The attention and suppression fields may drift away from the center, and only the suppression field can broaden (σ free). Our model recreated the experimental results shown in Fig. 3A (Fig. 4B, left) and Fig. 3D (Fig. 4B, right). The three colored crosses (magenta, black, green) highlighted example target locations that show increased, unchanged, and decreased firing rates due to attention, respectively. In each session, the standard deviation (or width) of the suppression field and the maximum allowable drift of the attention and suppression fields were fixed. Fig. 4C–E showed the histograms of 66 simulated sessions that matched two experimental criteria. One was the proportion of attention–suppressed locations (points below the diagonal in (B), left) lay between 10 % and 30 % (experiment: 21 %). The other one (Supplementary Fig. 3A) was the ratio of inside-RF locations (scaled passive rate 0.7–1.0 in (B), right) to outside-RF locations (0.2–0.5) exceeded 40 % (experiment: 62 %).

**Figure 4.**
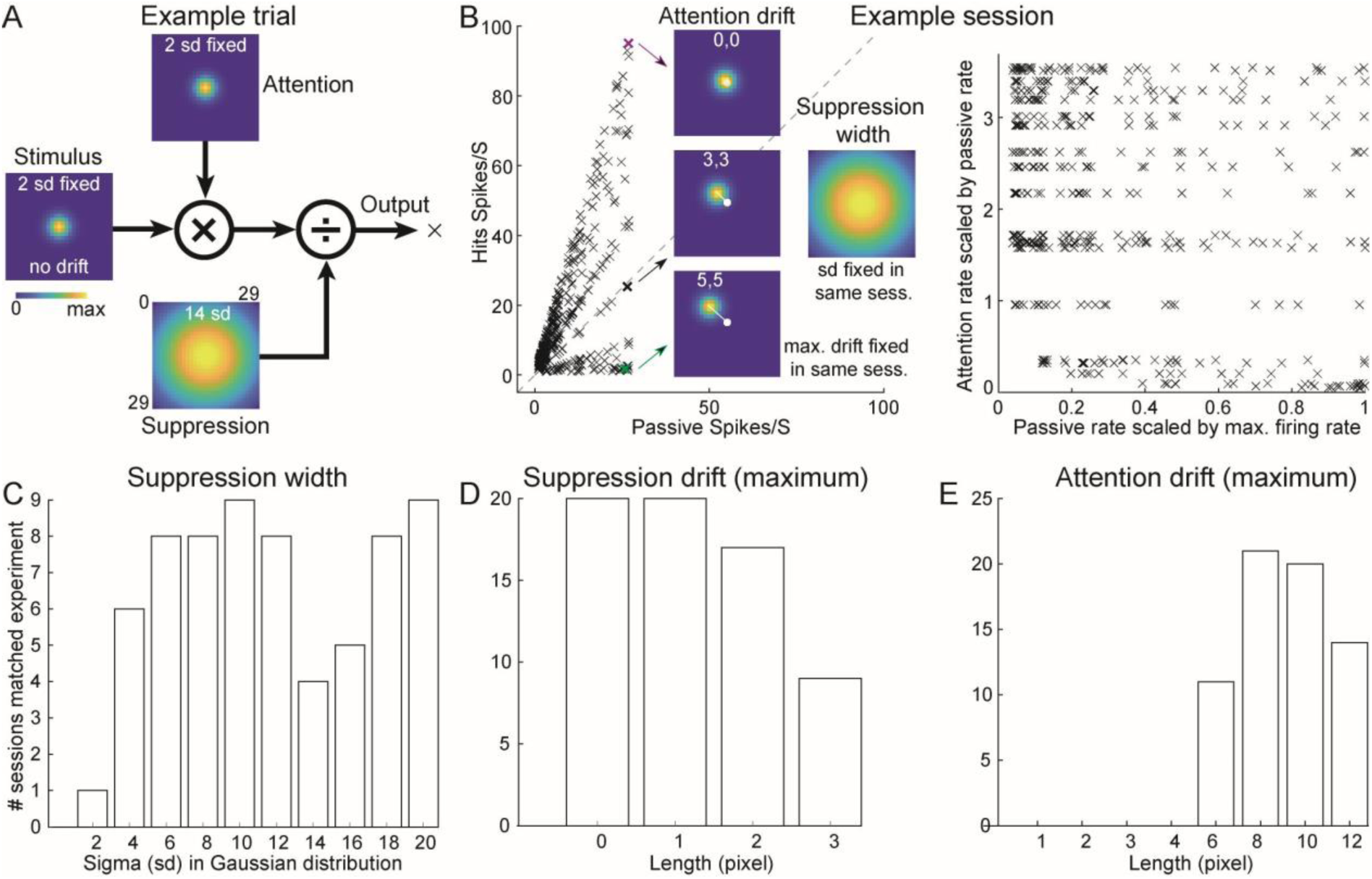
A normalization model reproduced experimental findings and revealed a candidate neural mechanism. (A) Three fields, multiplication, and division, in a normalization model. (B) Left, firing rates under behaving and passive conditions. Three example target locations have the same and fixed suppression width, but with different attention drift. (C) Suppression–field width (σ). (D) Maximum suppression–field drift. (E) Maximum allowable attention-field drift.

In all modeling sessions that matched experimental results, we found that the width of the suppression field has to be larger than the width of attention and stimulus fields (Fig. 4C). There was only one session succeeded with the same width as attention/stimulus fields, whereas many sessions succeeded even with a very broad width of suppression field. The remaining two parameters were the maximum allowable drift of suppression and attention drift. Most successful sessions had little or no drift in the suppression field (Fig. 4D). The allowed attention drift had to be larger than 6 pixels. To align with the experimental criteria (at least spike to at least one location), we excluded all trials (or locations) with output firing rates below one spike per second. All the retained locations ended up having actual attention drifts between 0 and 5 pixels (Supplementary Fig. 3B-C). Together, a normalization model recaptured our experimental observations. It further predicted that a widespread suppression in the spatial RFs played a necessary role.

### Widespread suppression is more prominent outside spatial RFs than outside spectral RFs

Our computational model suggests that the widespread suppression in the RFs contributes to the firing rate increase both inside and outside the spatial RFs. A suppressed firing rate relative to the spontaneous rate in response to nonpreferred locations was observed in the auditory cortex of passive listening cats (Mickey and Middlebrooks, 2003) and macaque (Woods et al., 2006). Here, we showed and reanalyzed the spatial RFs published previously (Remington and Wang, 2019) during the passive listening state (including marmosets used in this study). Figure 5A shows an example unit with a high spontaneous firing rate (20 spikes/s) and was suppressed at 18 sound locations (blue regions in the RF). Figure 5B further showed that the suppression was not limited to the ipsilateral space. Instead, it was widespread on both contralateral and ipsilateral spaces. To quantify suppression among the population (Fig. 5C, left), we choose 123 units and used the similar analysis we did previously for the spectral tunings (Supplementary Fig. 4A, B; Sadagopan and Wang, 2010, their Fig. 4). We also choose 59 units (among the same units used for spatial tunings) following the same criteria as the spatial tunings (Fig. 5C, right). Compared to the spectral RFs, both the suppressed area (7.6 vs 1.3) and stimuli (59 vs 28) were much larger in the spatial RFs. The small and narrow suppression in the spectral RFs was consistent with previous studies (onset and onset plus sustained responses in Sadagopan and Wang, 2010).

**Figure 5.**
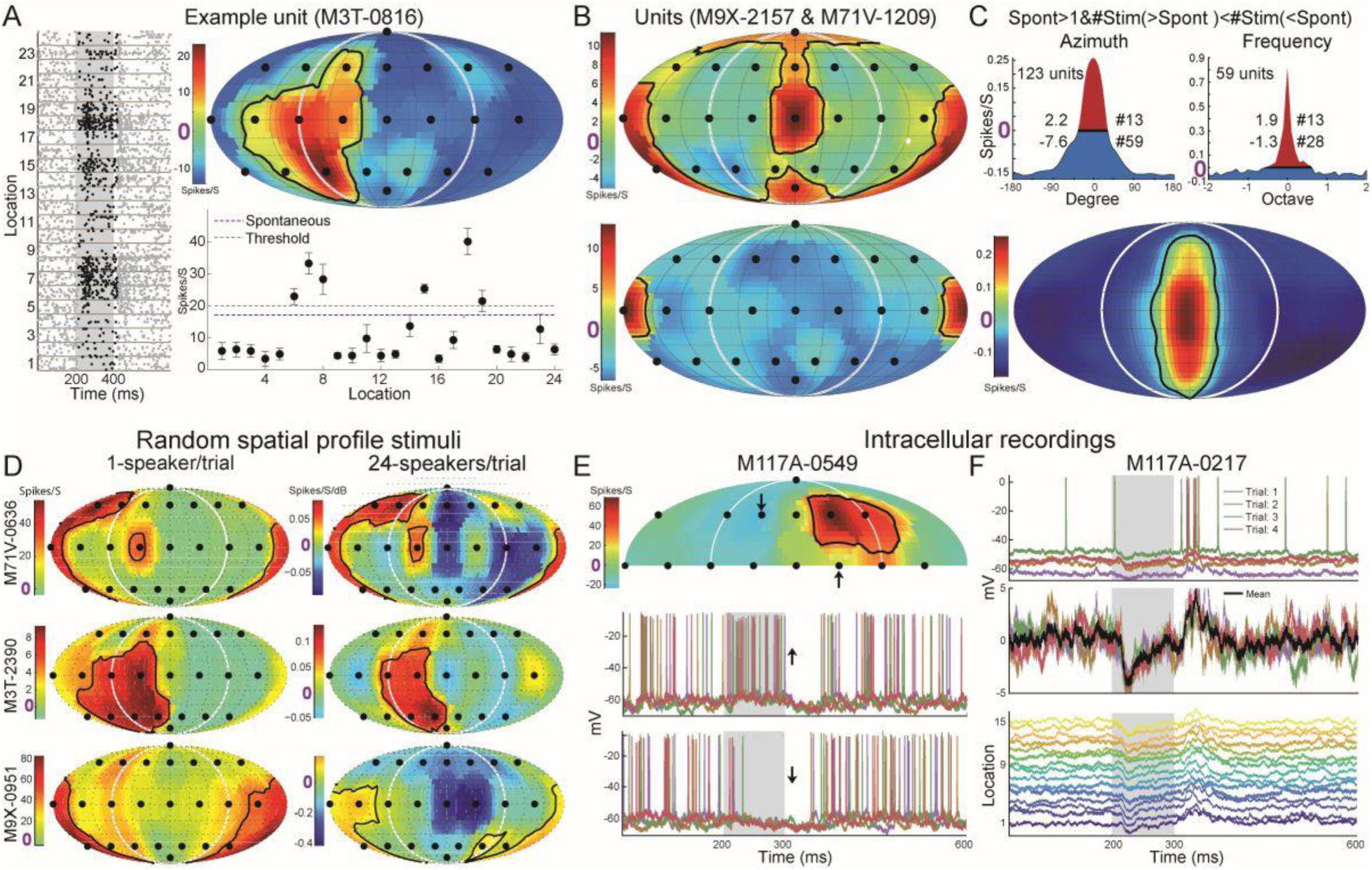
Firing rate suppression and membrane potential hyperpolarization evoked by stimuli from locations outside spatial RFs. (A) The spike raster (left), spatial RF (right top), and average firing and spontaneous rates (right bottom) of an example unit that preferred the contralateral bottom location. (B) Two example units that preferred sound from the meridian plane (top) and back (bottom) also showed suppressed firing rate to two or the majority of the locations (blue). (C) Widespread suppression among the population for spatial and spectral RFs. Top, units with larger than 1 spike/s spontaneous firing rate and larger number of suppressed stimuli (blue) than excited stimuli (red) were selected. The 1-dimensional azimuth (left) and frequency (right) tunings were normalized, circularly shifted, and averaged. Values on the left and right sides of each curve indicated the summed areas and stimuli number, respectively. Bottom, the 2-dimensional averaged spatial RFs. (D) The spatial RFs of three example units (rows) under two stimulus paradigms (columns). (E) Intracellular recordings from auditory cortex of an awake marmoset. Top, spatial RF calculated using the suprathreshold spikes. Bottom, membrane potentials at two locations that showed increased and decreased firing rate and membrane potential, respectively. (F) Top, membrane potential at one sound location across four repeats. Middle, raw, and averaged membrane potential with spikes subtracted. Bottom, mean membrane potential across all fifteen sound locations.

To observe firing rate suppression below the spontaneous firing rate shown in Fig. 5A-C, the neuron itself must exhibit sufficient spontaneous spikes. To drive neurons with low or no spontaneous firing (Fig. 5D, left) and measure the spatial RFs simultaneously, we designed a novel stimulus paradigm of “random spatial profile stimuli” that simultaneously deliver white noises from the array of 24 speakers with randomly chosen sound levels. This stimulus design was inspired by the random spectrum stimuli (RSS) (Barbour and Wang, 2003). RSS is a class of parametric wideband stimuli capable of driving neurons in multiple cortical fields and has also been applied to study auditory spatial tunings (Slee and Young, 2013) and spatial attention tasks (Chen et al., 2023). With these novel stimuli, we were able to reveal widespread suppression from off-RF locations that showed near-zero firing rates (Fig. 5D, right; Supplementary Fig. 5). Firing rate could only reveal neural responses above the spiking threshold; thus, the single unit recordings cannot reveal the suppressive effects below the threshold. Therefore, we used the intracellular recordings to measure the membrane potentials at different locations in the half spatial field from the passive listening marmosets (Gao et al., 2016). Figure 5E shows the spatial RF (top) and membrane potentials (bottom) at two different locations. Notice the membrane potentials were hyperpolarized (i.e., more negative) after (up arrow) and during (down arrow) sound stimuli. In the other example unit, there was no sound-evoked firing rate since its membrane potentials were suppressed at all fifteen sound locations. Together with novel sound stimuli and intracellular recordings, we were able to reveal widespread suppressions from many “unresponsive” sound locations.

## Discussion

We studied single-unit responses in the auditory cortex of marmosets while they performed a spatial discrimination task in different regions of the full spatial field. Comparing these responses to those measured while marmosets listened passively, a subset of neurons was observed to have increased firing rates to one or more target locations during task engagement. Effects at background locations were mixed. As the task involved a specific stimulus order which was different than that used to measure spatial receptive fields, we measured responses in an additional passive control condition in which the stimulus order was identical to that of the behavior condition. Effects were similar, even slightly larger, when comparing rates with the control condition, indicating that increased firing rates were not driven by stimulus order effects in most neurons. Increases occurred both within and outside of the classical spatial receptive field (typically defined as half-maximal firing rate area). Comparing effects of behavior between rostral (R/RT), caudal (CM/CL), and primary (A1) auditory areas, the largest effects were observed in CM/CL and the smallest in R/RT.

The observation that firing rate increases occurred at some and not all locations, while also distributed throughout RFs, indicates that effects in this task were stimulus specific. This has been observed previously in one study of spatial behavior (Benson et al., 1981), in which subjects were required to localize a sound source. In others, effects tended to occur throughout (Scott et al., 2007) or either in preferred or non-preferred portions of receptive fields (Benson and Hienz, 1978; Lee and Middlebrooks, 2011), indicating neuron-specific effects. Several studies in behaving ferrets have shown consistent stimulus-specific effects on frequency tuning, but not neuron-specific effects (Fritz et al., 2003, 2007). This dichotomy has also been observed in studies of spatial attention in the visual system, with neuron-specific effects including, but not limited to, multiplicative gain (Treue and Martínez Trujillo, 1999) and receptive field sharpening (Spitzer et al., 1988), and stimulus-specific effects exemplified by receptive field shifts (Womelsdorf et al., 2006). Another dichotomy in observations of the effects of behavioral engagement on firing rates in auditory cortex is that some studies have observed increased responses, generally for target stimuli (Benson and Hienz, 1978; Benson et al., 1981), while other studies have shown decreases, primarily to background stimuli (Otazu et al., 2009; Lee and Middlebrooks, 2011). Although differences have been observed due to task structure, such as between appetitive and aversive tasks (David et al., 2012), these differences were all in studies using positive reinforcement. One study was similar to ours in structure yet had dissimilar results: in cats performing a sound elevation discrimination task, responses were often decreased at non-preferred locations, while effects at preferred locations were mixed (Lee and Middlebrooks, 2011). It may be the case that behavior effects are dependent on specific details of task structure, such as the distributed backgrounds in Lee & Middlebrooks (2011) vs. the distributed targets here. Another possibility is that there exist species-specific differences between behavioral effects in different auditory areas. An experiment measuring responses in the same task in multiple auditory cortical areas showed increased responses in the posterior auditory field (Lee and Middlebrooks, 2013).

While firing rate increases could occur throughout receptive fields, firing rate increases occurred more consistently and strongly at target locations, and increases were likely to be larger where target/background contrast was positive in the passive condition. This suggests that these effects acted to tailor spatial representation for the specific behavior task. Previous studies of mammals performing sound location tasks have observed similar seemingly optimizing changes in spatial tuning. In the first, neural responses were recorded in macaques performing a dichotic listening task in which they were instructed to respond to sounds played to the left or right ear. In this task, a population of contralateral preferring neurons responded more strongly to contralateral locations when the target was contralateral (Benson and Hienz, 1978). In addition, cats trained to discriminate sound elevation along the complete azimuthal dimension displayed depressed responses at non-preferred locations (Lee and Middlebrooks, 2011). In both of these cases, changes occurred that were optimized to the task in the context of the underlying spatial (or binaural) tuning. It has been suggested that this is an underlying high-level behavior of spatial processing in auditory cortex (Lee and Middlebrooks, 2011); our results seem to add support to this hypothesis.

Several studies have shown that the distributions of spatial tuning properties vary quantitatively (but not qualitatively) between auditory areas along the rostral-caudal axis, with neurons in caudal areas, statistically, displaying higher selectivity for spatial locations than those in rostral areas and primary auditory cortex (Tian et al., 2001; Stecker et al., 2003; Woods et al., 2006; Lee and Middlebrooks, 2013; Remington and Wang, 2019). Differences then might be expected in the way tasks involving sound location affect caudal vs. primary and rostral auditory cortex. A study comparing effects between A1, PAF, and DZ in cats found a lower fraction of neurons in the posterior auditory field and dorsal zone, which sharpened tuning compared with A1 (Lee and Middlebrooks, 2013). Additionally, neurons in the posterior auditory field were the only population to consistently increase their responses during behavior. This report represents the first time behavior effects on spatial responses in primates have been directly compared between multiple auditory areas, including significant recordings from caudal areas, although effects of behavior outside of the primary auditory cortex have been measured previously, primarily in rostral and lateral areas (Benson and Hienz, 1978; Benson et al., 1981). For the most part, quantitative, not qualitative difference in effects between areas mirrors quantitative, but not qualitative differences in tuning properties between areas (Stecker et al., 2003; Woods et al., 2006; Zhou and Wang, 2012), although the present data do suggest the possibility of multiplicative in addition to additive gain in areas CM/CL. The most dramatic difference in effect size was the apparent lack of consistent behavior effects in R/RT; however, it is possible that these differences were the result of stimulus order. When behaving firing rates in R/RT were compared with the control condition with identical stimulus order, increases became apparent, albeit still smaller than in A1 and CM/CL. To our knowledge, no study has specifically compared the effects of stimulus interaction over long time scales across the rostral-caudal axis.

The varied but significant effects of behavioral context on spatial receptive fields in auditory cortex suggest that spatial tuning in the passive condition provides only part of the picture of spatial representation in auditory cortex. These and other behavior studies may reconcile disparate observations regarding spatial tuning in auditory cortex. For example, studies of spatial processing in anesthetized animals have observed a preponderance of neurons with very broad spatial receptive fields, which almost universally increase in size with increasing sound level (Brugge et al., 1994, 1996; Mrsic-Flogel et al., 2005). However, in awake animals, spatial receptive fields tend to be smaller and do not uniformly increase in size with sound level (Mickey and Middlebrooks, 2003; Woods et al., 2006; Zhou and Wang, 2012; Remington and Wang, 2019). The observation that large firing rate increases can occur far from the best location may be evidence of broader inputs that are masked in the passive awake condition when compared to the anesthetized state. Conversely, another behavior task may lead to still more selective tuning compared to the passive state (Lee and Middlebrooks, 2011). We therefore believe that the true nature of spatial representation may not be well understood by studying non-behavioral subjects, and that a complete picture will require further behavior studies.

## Methods

### Animal preparation and electrophysiological procedures

Experimental procedures were approved by the Institutional Animal Care and Use Committee of the Johns Hopkins University following National Institutes of Health guidelines. A chronic recording preparation was used to record single-neuron activity in the auditory cortex (left hemisphere) of two female common marmoset monkeys (Callithrix jacchus). Both subjects were trained to sit in a custom-designed primate chair and perform a simple auditory detection task (Remington et al., 2012) and then a spatial discrimination task (Chen et al., 2023). After training, two stainless steel headposts were attached to the skull under sterile conditions with the animal deeply anesthetized by isoflurane (0.5–2.0%, mixed with 50% O2 and 50% nitrous oxide). The headposts served to maintain a stable head orientation of the subject during electrophysiological recordings, although only one post was fixed in this study. To access the auditory cortex, small craniotomies (1.0 or 1.1 mm in diameter) were made in the skull over the superior temporal gyrus to allow for penetration of single electrodes (tungsten electrodes, 2- to 5-MΩ impedance, A-M Systems, Carlsborg, WA) into the brain via a hydraulic microdrive (Trent-Wells, Los Angeles, CA). Single-unit activity was sorted online using template-based spike-sorting (MSD, Alpha Omega Engineering) and analyzed using custom programs written in Matlab (Mathworks, Natick, MA). We used intracellular recording procedures identical to those in Gao et al. (2016). Recordings were made in the auditory cortex through the intact dura with a concentric recording pipette and guide-tube assembly. Sharp recording pipettes were quartz glass pulled on a laser puller (P-2000, Sutter), and guide tubes were borosilicate glass pulled on a conventional puller (P-97, Sutter). The electrode assembly was advanced perpendicular to the cortical surface using a motorized micromanipulator (DMA-1510, Narishige). Electrical signals were amplified (Axoclamp 2B, Molecular Devices), digitized (RX6, Tucker-Davis Technologies), and saved with custom MATLAB code (MathWorks).

### Acoustic stimuli and receptive field characterization

Experiments were conducted in a double-walled sound-attenuating chamber (Industrial Acoustics, IAC, New York) with internal walls, ceiling, and floor lined with ∼3-inch acoustic absorption foam (Sonex). Acoustic stimuli were delivered using an array of 24 speakers (FT28D, Dome Tweeter, Fostex) covering a complete sphere. The loudspeakers were mounted at a distance of 1 m from an animal’s head and covered 5 Elevations (ELs) at 45° spacing and several Azimuths (AZs). One speaker was located directly above the animal, 7 speakers each were evenly spaced at ± 45° EL (AZ at –45° EL: ±25.7°, ±77.1°, ±128.6° and 180°; AZ at 45° EL: 0°, ±51.4°, ±102.9° and ±154.3°), 8 speakers were evenly positioned at 0° EL (AZ: 0°, ±45°, ±90°, ±135°, 180°), and finally 1 speaker was located at -67.5° EL at 0° AZ. Subjects sat in a wire mesh primate chair mounted onto a single stainless steel bar such that the animal’s head was centered in the room. Marmosets were head-fixed for all recordings. In this text, positive AZ angles correspond to speakers ipsilateral to the recording site or to an ipsilateral shift if changes in azimuth were analyzed. During experiments, eye position was not controlled.

Stimuli were generated in Matlab (Mathworks) at a sampling rate of 97.7 kHz using custom software. Digital signals were converted to analog (RX6, 2-channel D/A, Tucker-Davis Technologies), then analog signals were attenuated (PA5 x2, Tucker-Davis Technologies), power amplified (Crown Audio x2), and played through a chosen channel of a power multiplexer (PM2R x2, 16-channels, Tucker-Davis Technologies). Loudspeakers had a relatively flat frequency response curve (± 3-7 dB) and minimal spectral variation across speakers (< 7 dB re mean) across the range of frequencies of the stimuli used; all large (5-7 dB) spectral deviations occurred in narrow bandwidths near the upper limit of speakers’ frequency range (above 28 kHz), above the first spectral notch measured in marmoset head related transfer functions (Slee and Young, 2010). Neurons were characterized for frequency, intensity, and spatial tuning. For frequency tuning, stimuli consisted of pure tones, band-pass filtered Gaussian noise, Random Spectral Shape (RSS) stimuli (Yu and Young, 2000; Barbour and Wang, 2003), and occasionally frequency modulated (FM) sweeps. We sampled the frequency axis in 0.1 octave steps, typically over a 4-octave range (2-32 kHz). All firing rates were calculated over a time window beginning 15 ms after stimulus onset and 20ms after stimulus offset. Best frequency was defined as the frequency that led to the maximum evoked significant firing rate, or for neurons only driven by RSS stimuli, the highest calculated RSS weight. For spatial tuning, stimuli included band-pass filtered unfrozen Gaussian noise, single RSS stimulus tokens, and occasionally FM sweeps. All stimuli used to measure spatial receptive fields were either band-pass filtered or constructed to have energy between 2 and 32 kHz. When possible, spatial receptive fields were measured at multiple stimulus intensities. Stimuli were typically 200ms long with 10ms cosine ramps and delivered in pseudorandom order, and, except RSS stimulus sets, delivered between 5 and 10 times and averaged to generate tuning functions.

We characterized spatial receptive fields in a spatially dense acoustic environment by playing sets of broadband sounds from the entire 24-speaker array simultaneously, randomizing the sound level from each speaker. This stimulus delivery paradigm was adapted from a similar method (random spectral shape stimuli) used to study spectral processing in the auditory system (Yu and Young, 2000; Barbour and Wang, 2003). The complete set of stimuli comprised a stimulus matrix **Λ** of intensities in which the rows represent individual stimuli and the columns represent the individual speakers. An RSP set is constructed to sample the space of all possible 24-location spatial profiles in such a way that weighting functions can be calculated to describe the spatial tuning to spatially dense stimuli. To do this, the set must be constructed such that the levels of each speaker are statistically independent across all stimuli. This condition is satisfied if the location intensity autocorrelation matrix is equal to the identity matrix:

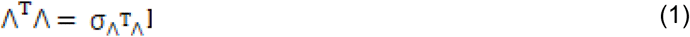

This constraint can only be met for a matrix having more rows than columns. Therefore, the minimum number of stimuli required to construct a linear estimate of the spatial weighting function in this case is 25. The linear spatial weighting function can be calculated from responses to an RSP set using the following equation:

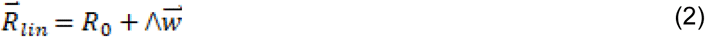

Where 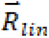 is a column vector of *m* rate values predicted in response to a set of *m* different RSP stimuli, *R*_0_ is the firing rate to an RSP stimulus with a flat spatial profile, Λ is the mean-adjusted intensity matrix, and 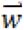 is the 24-value linear weighting vector. This equation is referred to as the linear synthesis equation. The weighting function is calculated as:

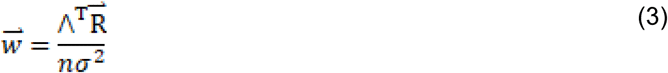

Where 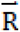 is the firing rate vector (in spikes/second) to the RSP stimulus set, *n* is the number of stimuli in the set, and σ^2^ is the variance of the sound levels at each speaker. Weights 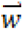 are expressed in units of spikes/second/dB. This equation is referred to as the analysis equation. Thus, when the intensity is increased at a location with a positive weight function, the firing rate should increase as well.

### Identification of A1, the rostral fields R/RT, and caudal areas CM/CL

In the marmoset, A1 is situated largely ventral to the lateral sulcus on the superior temporal plane and, similar to other primate species, exhibits a low-to-high topographical frequency gradient along the rostral-caudal axis. The boundary between A1 and the rostral field R can be identified by a downward-to-upward frequency gradient reversal along the rostral-caudal axis. Conversely, areas CL and CM can be identified by an abrupt decrease of best frequency at the high frequency (caudal) border of A1 (Kaas and Hackett, 2000; Song et al., 2022). In this and our previous studies (Remington and Wang, 2019), the boundaries between R/RT and A1 and A1 and CM/CL were set by plotting the average best frequency along the rostral-caudal axis, approximately parallel to the lateral sulcus, and setting a boundary at the local minimum and maximum, respectively, between the two areas. We did not separate neurons further into R/RT or CM/CL, although some studies have found differences in spatial selectivity between areas CM and CL in macaques (Woods et al., 2006; Kusmierek and Rauschecker, 2014).

### Spatial discrimination task

We chose to implement a Go/No-Go type task suited for discrimination behavior. Fig. 1A illustrates the behavior paradigm. The objective in a Go/No-Go task is to respond (a lick at the feeding tube) to target sounds to receive a food reward while withholding responses when a target is not presented. Here, each trial was composed of a variable length “intertrial interval” in which sounds were played only from background locations and a fixed length “response interval” during which target and background locations alternated. Intertrial interval length was randomized between approximately three and ten stimuli, and the response interval included four target/background alternations. Behavioral responses during the intertrial interval resulted in a time-out, and sometimes a puff of air to the base of the tail, followed by a restarting of the intertrial interval. After the intertrial interval ended, target stimuli were alternated with the background sounds during the response interval. Trials ended when the response interval expired or a lick was detected during the response interval. Behavioral responses during this time were reinforced with approximately 0.1– 0.2 ml of food reward. If no response was detected, the next intertrial interval began immediately. One third of trials were sham trials in which stimulus location did not change, and no reward was given for behavioral responses. False alarms were measured with sham trials.

Four target/background configurations were used. The background location was 45° lateral to the midline (front and back, contralateral and ipsilateral), and the target locations were the most lateral positions (±90°; same in all conditions), and also 45° above and below the horizon, but in the same azimuthal quadrant as the background location. For targets above and below the background location, their azimuth locations were either 51° or 25.5° lateral to the midline (one of each per condition). A diagram of one such condition is shown in Fig. 1B, C. Stimuli were 200ms in length, and the interstimulus interval was approximately 500ms, resulting in a stimulus onset asynchrony of approximately 700ms. Sound level was roved either ±5 dB SPL or ±10 dB SPL to prevent the use of changes in sound level as a perceptual cue. Mean sound level was chosen to be within the flattest portion of each neuron’s rate-level function.

### Effects of behavior on spatial responses

All comparisons between conditions were done using “driven” firing rates, equal to the raw firing rate minus the average spontaneous firing rate measured in that condition. We calculated three measures to quantify the difference in firing rate between behaving and passive conditions. Modulation index (MI) is a measure of the response difference at a particular location scaled by the combined response strength at that location:

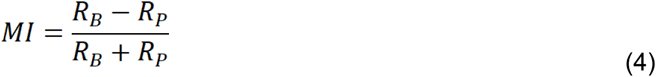

where *R_B_* is the firing rate in the behaving condition, and *R_P_* is the firing rate in the passive condition. The hit rate increase (Y-axis in Fig. 3D-F) is a measure of the response increase at a location relative to the maximum passive firing rate:

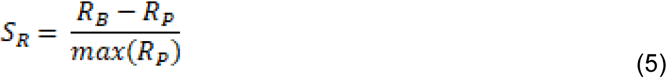

Where *max*(*R_P_*) is the maximum firing rate among all locations in the passive spatial RFs.

We compared rates during active behavior to two different passive conditions. The first condition is the standard spatial receptive field measured by playing sounds from all 24 speaker locations in a randomized order. To control for effects of stimulus order, which can include suppression or facilitation of neural responses (Chen et al., 2025), we compared firing rates in the behavior condition to a second condition in which stimulus delivery was identical to the behavior condition except that stimuli did not stop if the animal responded. Data were rejected if a response was made during this control condition, although this was rarely the case. Subjects were cued to the beginning of behavior sessions by alternating the house light on and off.

### Statistics and modeling

Wilcoxon rank-sum tests were used to evaluate the population medians when evaluating statistical significance of populations of values, except when testing for deviations from zero mean, in which case t-tests were used. Correlation analyses were based on Spearman’s correlation coefficient. All data analyzed in this study were for neurons and locations that displayed a driven firing rate (p < .001, minimum one spike per stimulus presentation) in either the behaving or passive condition. Bonferroni–adjusted p-values are reported for tests with multiple pairwise comparisons. Our normalization model was inspired by two previous studies (Reynolds and Heeger, 2009; Ohshiro et al., 2011). The first paper proposed a classical one-dimensional normalization model:

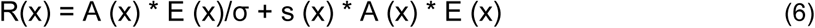

Which included R (neuron firing rate), x (center of receptive field, Gaussian distribution), A (attention field), E (stimulus field), (σ semi-saturation constant), and S (suppression field). The second paper proposed a two-dimensional normalization model for multisensory integration (we only used one modality) without an attention field:

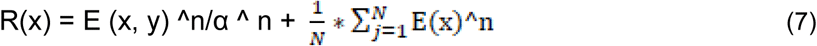

Where n represents the exponent of the output nonlinearity.

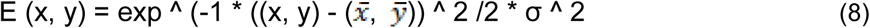

Where 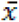 and 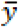 represent the center of the receptive field. The standard deviation (s.d.) or the width of the receptive field was determined by σ. The key difference from the previous model was the normalization part, which was a weighted sum over nearby neurons. Our two-dimensional normalization model was:

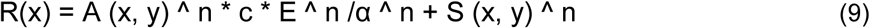

Here, *n* was fixed at 2 and α at 32. The size of the receptive field was 29 × 29, and the center of the stimulus field *E* was fixed at the center (15, 15). The centers of the attention field *A* and suppression field *S* were allowed to drift from this center. The random intensity of sensory inputs was controlled by *c*, which ranged from 8.75 + [1, 2, 3, 4, 5]. The maximum allowable drift of the attention and suppression fields was [1, 2, 3, 4, 6, 8, 10, 12] and [0, 1, 2, 3], respectively. The size of the receptive field σ was fixed for the attention and stimulus fields but ranged between [2, 4, 6, 8, 10, 12, 14, 16, 18, 20]. For the passive condition without attention, we removed the attention field A *(x, y)*.

## Conflict of interest

The authors declare no competing financial interests.

## Acknowledgements

We thank J. Estes and N. Sotuyo for assistance with animal care and M. Osmanski for feedback on behavioral training and the manuscript. The intracellular recording examples were obtained by Dr. Yunyan Wang while a postdoctoral fellow in the Wang Lab. Support was contributed by US National Institutes of Health grant DC003180 (X.W.).

## Supplementary Figures

**Supplementary Figure 1.**
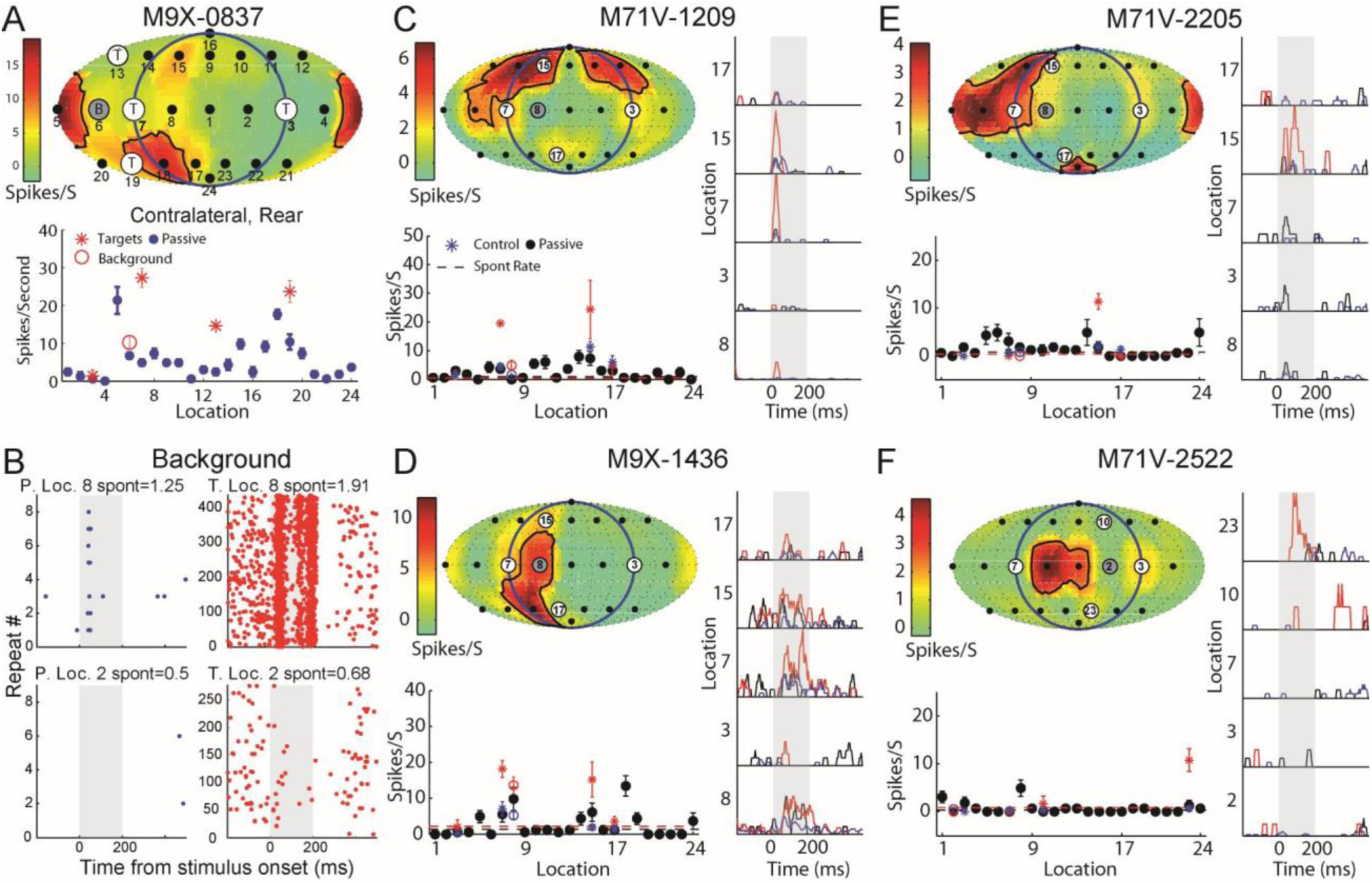
More example units with firing rate increase outside the spatial RFs (related to Figure 2). (A) An example session (same unit in Fig. 2A) where the background location (#6) and two target locations (#13 and #19) were located at the rear of the animal. (B) Spike raster from two example units (Fig. 2A and 2F) at the background locations (#8 and #2) under passive (left) and target (right) conditions. Notice there were many more trials under the target locations. (C-F) Four more example units. Here, we used black, blue, and red colors to represent 24 locations during passive, control, and behaving conditions (five targets/background locations), respectively. Spontaneous firing rates were indicated with colored dashed lines.

**Supplementary Figure 2.**
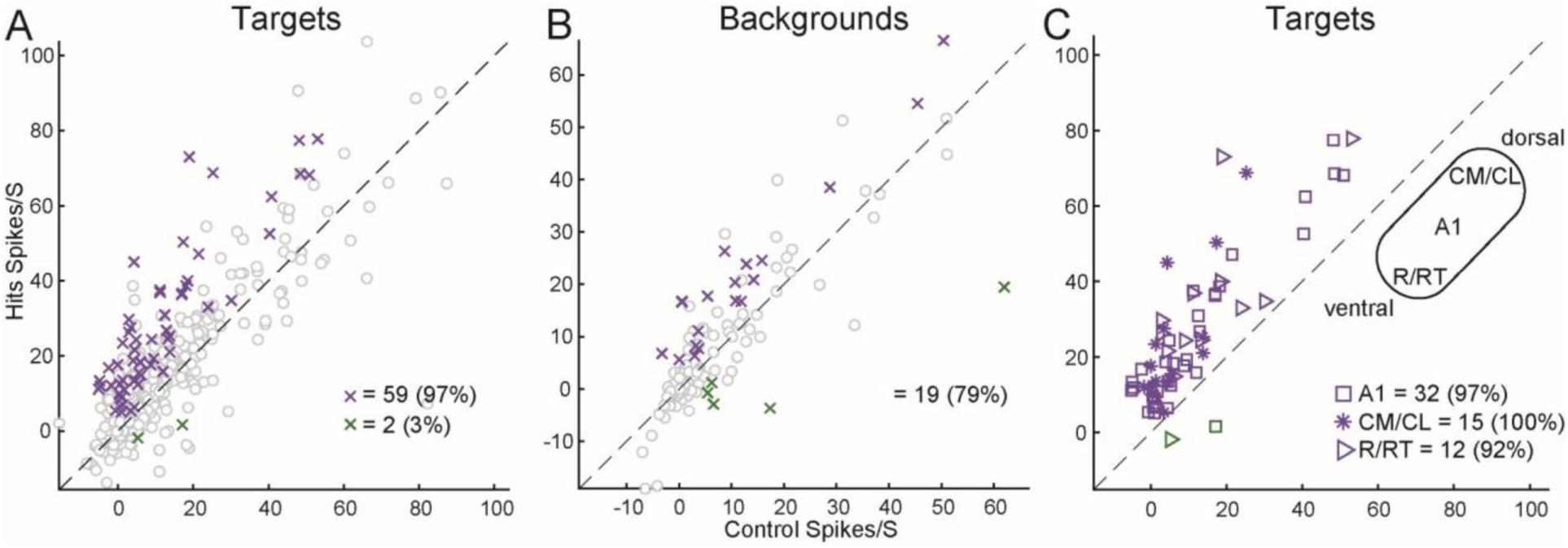
Firing rate during the behavior and control conditions (related to Figure 3A-C). (A) Comparisons of firing rates in the hits (i.e., successful behavioral choice) and control conditions at all target locations. Gray circles represent the nonsignificantly modulated target locations (249 above and 199 below the diagonal). (B) Similar to (A) but at the background locations. There were 53 and 55 nonsignificant background locations above and below the diagonal, respectively. (C) Same data as the significantly increased (pink) and decreased (green) firing rate at target locations shown in (A), but with three different shapes (asterisk, square, and right-pointing triangle) to distinguish the three auditory cortical areas (A1, CM/CL, and R/RT).

**Supplementary Figure 3.**
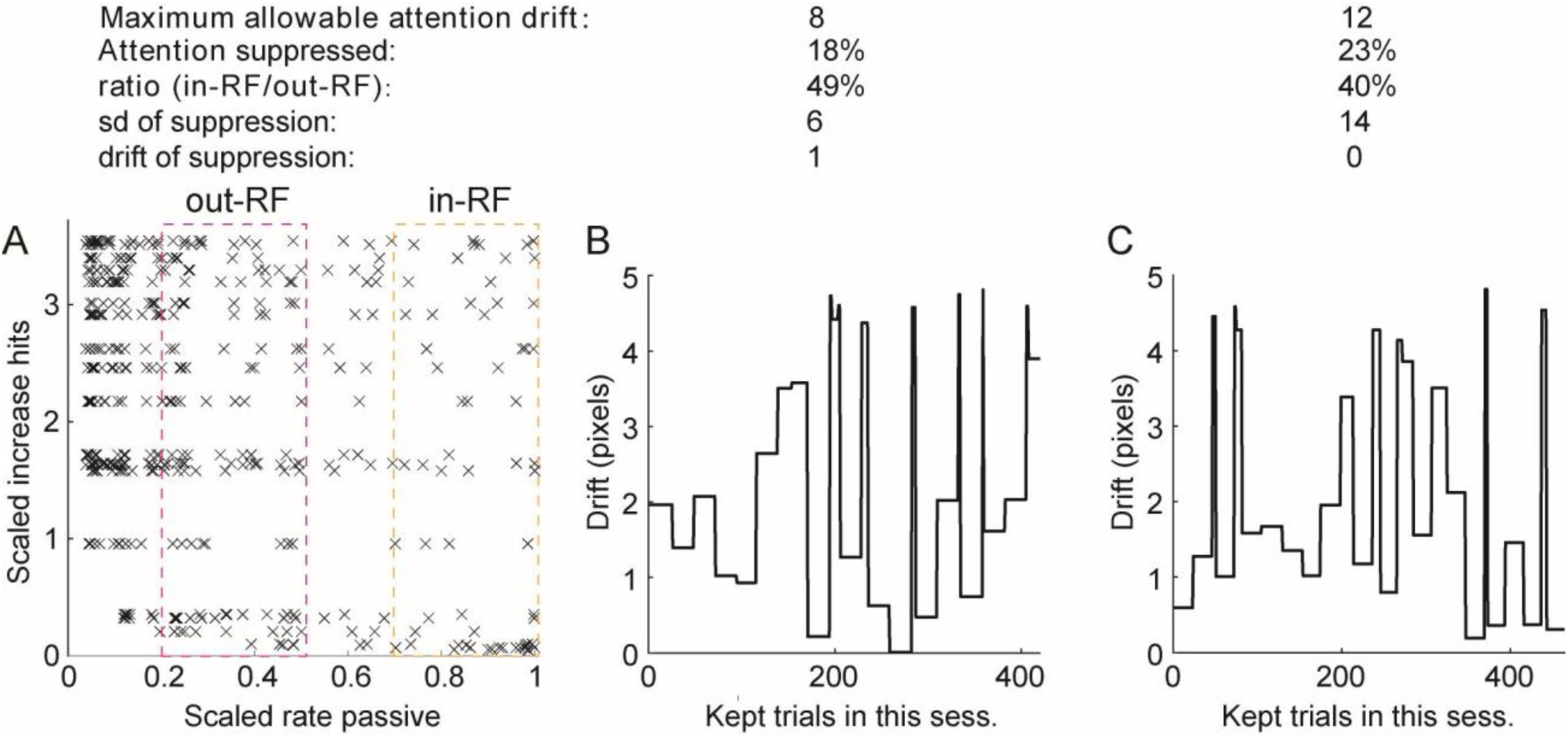
Increased firing rates to target locations increased firing rates relative to background locations (related to Figure 4). (A) Firing rate during task plotted vs. passive firing rate for all data points that were larger than 1, both scaled by the maximum firing rate in the passive condition. The scaled passive rates between 0.2 to 0.5 and 0.7 to 1 were considered as outside and inside of receptive fields, respectively. (B-C) The drift of attention fields in all kept trials from two example sessions. The maximum drifts were only 5 pixels in the kept trials, although the allowable drifts were much larger.

**Supplementary Figure 4.**
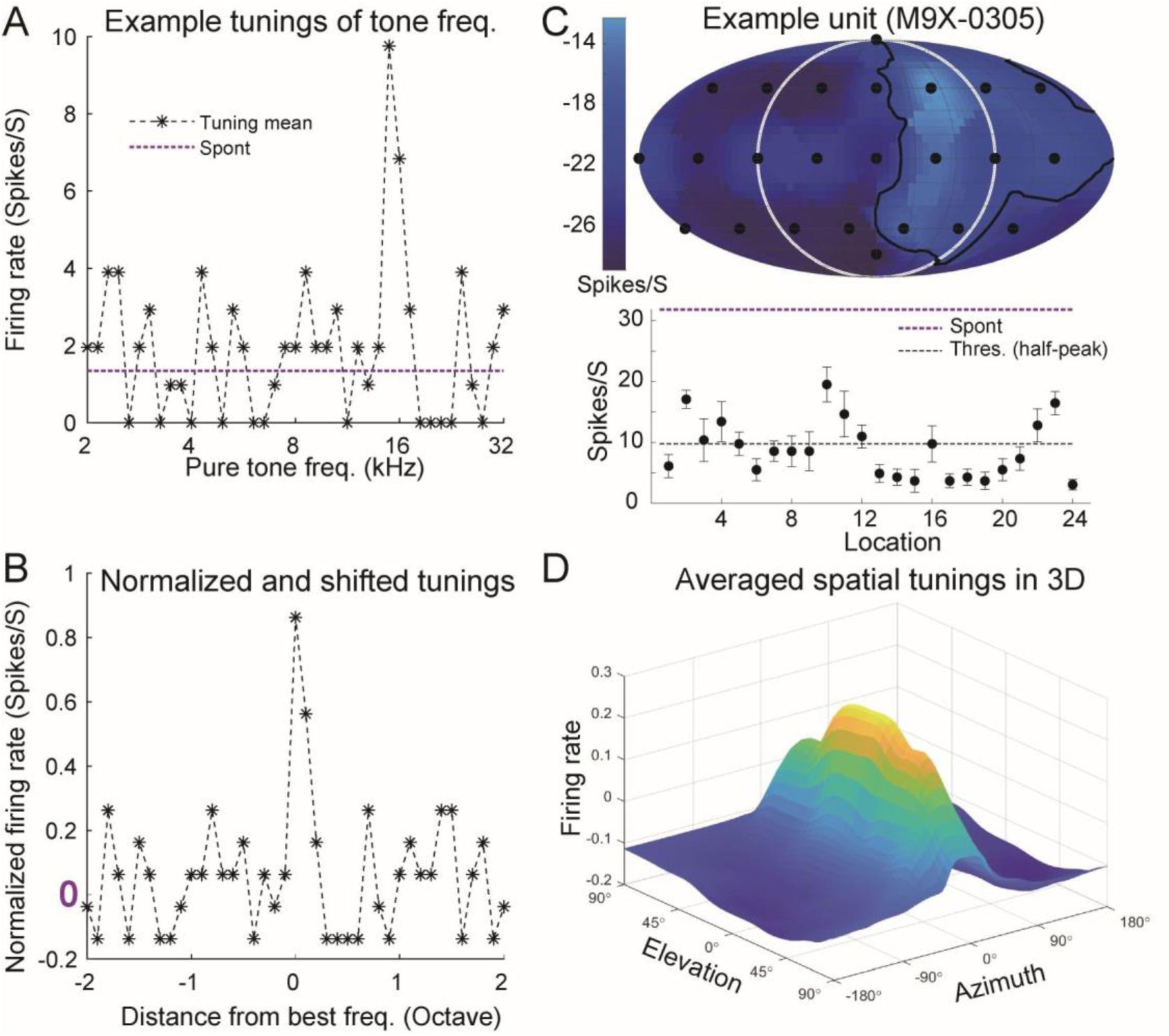
Suppressed neural firing rate to the nonpreferred sound stimuli (related to Figure 5C). (A) Averaged neural firing rate to 31 sound frequencies (4 octaves, 8 stimuli per octave) of an example unit. (B) The tuning curve was normalized by the maximum firing rate at 16 kHz (the spontaneous firing rate equated to “0”). It was further circularly shifted so that there were 15 frequencies (2 octaves) at both the left and right sides of the peak firing rate. (C) An example unit that was suppressed at all 24 sound locations. Notice that the neuron was still significantly tuned to sound locations (ANOVA, p<0.01). (D) A 3D view of averaged spatial receptive fields with both color and height at the third axis represented the firing rate.

**Supplementary Figure 5.**
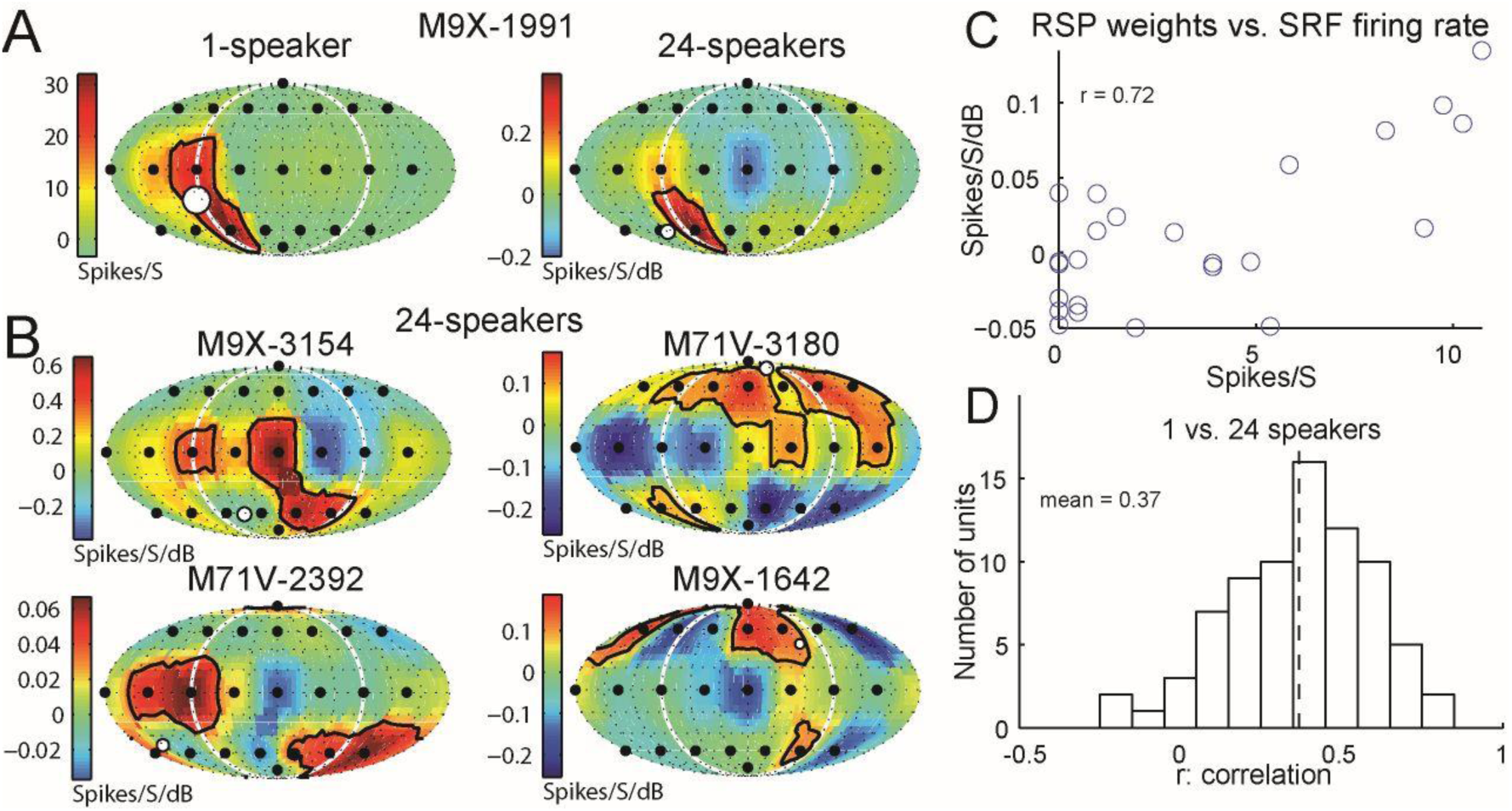
Suppressed neural firing rate to the random spatial profile (RSP) sound stimuli (related to Figure 5D). (A) An example unit showed consistent spatial receptive fields under two stimulus paradigms. The position and size of white dots indicate the center of receptive fields and their tuning selectivity, respectively. (B) Four more example units all showed suppressed firing rates at more than one sound location using the RSP stimuli. (C) The scatter plot (X-axis: 1-speaker, Y-axis: 24-speakers) of neural activities at 24 sound locations under two stimulus paradigms. The correlation (r) of neural activities was 0.72. (D) The histogram of correlation for all 77 units.

## Notes

https://github.com/ccg1988/SpatialAttention

### Competing Interest Statement

The authors have declared no competing interest.

